# GRIDGENE: Guided Region Identification based on Density of GENEs – a transcript density-based approach to characterize tissues by spatial transcriptomics

**DOI:** 10.1101/2025.08.14.670318

**Authors:** A. M. Sequeira, M. E. Ijsselsteijn, M. Rocha, J. Roelands, Noel F.C.C. de Miranda

## Abstract

Spatial omics brought unprecedented power to study biological processes within tissues while preserving spatial context and morphology. Most spatial proteomics and transcriptomics analyses methods are cell-centric, relying on cell segmentation to identify and characterize individual cells before downstream tasks. However, certain biological questions may be better addressed using cell-free approaches, which also eliminate unnecessary computations when cell segmentation is not essential. To address this need, we developed GRIDGENE (Guided Region Identification based on Density of GENEs), an approach for defining regions of interest based on transcript density. GRIDGENE enables the identification of biologically relevant tissue compartments, including interfaces between regions, phenotype-enriched areas, and zones defined by specific gene signatures, supporting analyses such as pathway enrichment. We demonstrated the utility of GRIDGENE by applying it to spatial transcriptomics data from CosMx and Xenium platforms in colorectal cancer (CRC) samples.

By bypassing cell segmentation, our approach enables flexible analysis of spatial omics data, supporting the study of biological processes across diverse tissue structures and microenvironments. Nevertheless, GRIDGENE can be easily integrated with cell segmentation strategies for complementary analyses. GRIDGENE thus broadens the analytical toolkit for spatial omics, enabling both cell-free and cell-based insights.

## 1. Introduction

Spatial omics has revolutionized the study of tissue microenvironments, providing unprecedented insight into spatial organization, cellular heterogeneity, and cell-to-cell interactions (1–3). In particular, for spatial transcriptomics (ST), several commercially available technologies enable the detection of hundreds of RNA transcripts at subcellular resolution (4,5), significantly expanding our ability to explore complex biological systems. However, these advancements also expose limitations in current analytic approaches.

Standard analysis pipelines for single-molecule ST data are primarily cell-centric, relying on image segmentation to identify and classify cells before downstream analysis (4–7). Tissues comprise heterogeneous cell populations with diverse shapes, sizes, and packing densities, often lacking a consistent orientation, which makes accurate segmentation inherently challenging and increases the risk of information loss. These complexities of spatial data can limit the effectiveness of cell-focused approaches, potentially hindering the detection of specific cell populations. Furthermore, cell segmentation and characterization may not always be necessary for understanding biological processes in spatial contexts.

An alternative to cell-centric methods is the use of segmentation-free approaches, which analyse spatial omics data without relying on cell segmentation. Instead, these methods define spatial neighbourhoods or compartments based on the underlying distribution of molecular points. Studies have shown that segmentation-free approaches can accurately identify cells (8,9). Additionally, there has been a recent surge in methods designed for unsupervised identification of spatial domains (10,11) and transcript expression patterns (12,13). For example, Points2Regions is a Python tool that identifies regions based on mRNA composition within spatial bins of varying sizes (14).

While unsupervised methods are valuable, certain studies may benefit from a more targeted approach. Researchers often need to define specific regions based on prior knowledge to address focused questions, such as identifying tissue compartments with distinct cellular compositions (e.g., cancer vs. stroma), analysing biological processes occurring at specific sites within a tissue, or characterizing regions dominated by particular cell types or gene expression profiles. Moreover, rare cell types or those defined by only a few transcripts may be overlooked during segmentation or clustering. Directly identifying regions of interest based on transcript density offers a straightforward and effective solution to these challenges.

This paper introduces GRIDGENE (Guided Region Identification based on Density of GENEs), a Python package for supervised exploration of spatial omics data. GRIDGENE allows researchers to identify regions of interest by incorporating prior biological knowledge such as expression profiles of specific tissue compartments, cell types, or annotated biological pathways. Using high-resolution single-molecule ST data from CosMx (5) and Xenium (4) on colorectal cancer (CRC) samples, this study demonstrates how GRIDGENE can extract insights without relying on traditional cell segmentation pipelines. GRIDGENE’s flexibility makes it a powerful tool for both exploratory and hypothesis-driven spatial-omics research.

## 2. Results

### 2.1 Description of GRIDGENE

GRIDGENE (Guided Region Identification based on Density of GENEs) is a Python package designed for identifying and analysing regions of interest based on transcript density, enabling rapid and customizable exploration of tissues in spatial omics data. It allows users to create tissue masks based on prior knowledge while being a flexible tool. The package is freely available at https://github.com/deMirandaLab/GRIDGENE.

GRIDGENE facilitates the analysis of ST data by providing a step-by-step workflow for identifying and characterizing regions of interest (Figure 1). First, users define areas of interest within tissues based on the density of specific transcripts (*Contour identification*, Figure 1A), which GRIDGENE converts into digital masks (*Mask generation*, Figure 1B). Users can expand these masks, for example, to enable specific analyses of areas surrounding a tissue compartment. GRIDGENE can retrieve relevant data within these masks, including transcript counts or morphological features, while preserving the topological and/or hierarchical relationship between masks (Figure 1C). Finally, the generated masks can be overlaid onto existing cell segmentation data, enabling rapid annotation of cells based on their localization within defined tissue compartments (Figure 1D). This integration not only helps identify which cells underlie specific gene expression signatures but also reveals their spatial distribution across compartments and allows quantification of cell densities within each region. This provides a complementary approach to traditional cell segmentation, combining transcript density information with cell-level annotation. Each submodule of GRIDGENE is explained in detail below.

**Figure 1.**
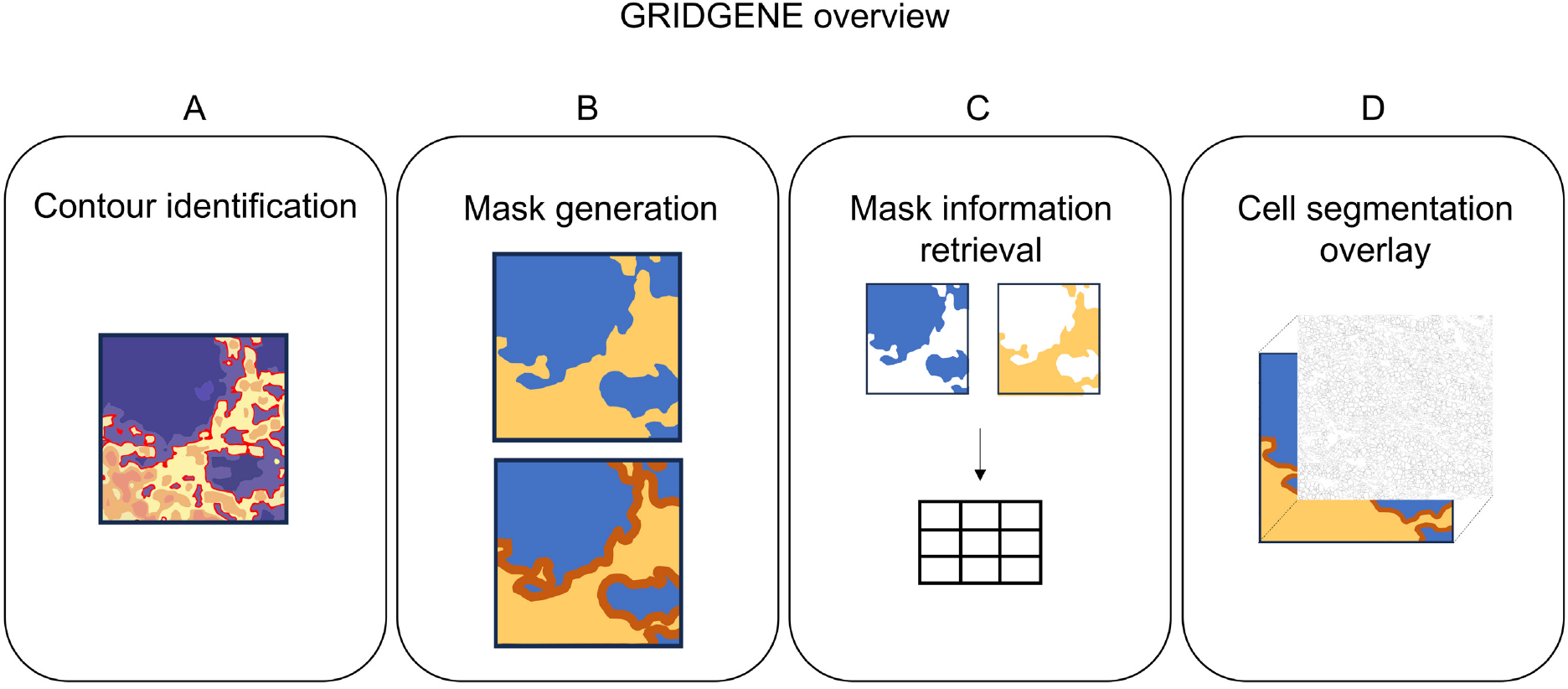
GRIDGENE overview. The framework is composed of four main steps: Contour identification, where regions of interest are delimited based on transcript density, neighbourhood counting, or SOM based; Mask generation, where the identified contours are converted into binary masks that can be further modified through morphological operations or expanded; Mask information retrieval, where transcript counts, morphological descriptors and hierarchical relationships between masks are extracted; and Cell segmentation overlay, where the generated masks are integrated with single cell segmentation data.

#### Contour identification

GRIDGENE’s primary objective is to identify and outline areas of interest—contouring. It accomplishes this by detecting regions with a high density of user-defined transcripts and generating contours around them (Figure 2). GRIDGENE primarily relies on the convolutional sum method to locate these regions. In the convolutional sum method (Figure 2A), a sliding window moves across the tissue, and at each position, it sums the number of transcripts within the window. This sum is then assigned to the central point of the window, creating a map of local transcript density across the tissue. Contours are then defined using specific rules, such as highlighting regions where values exceed or fall below a given transcript count threshold.

**Figure 2.**
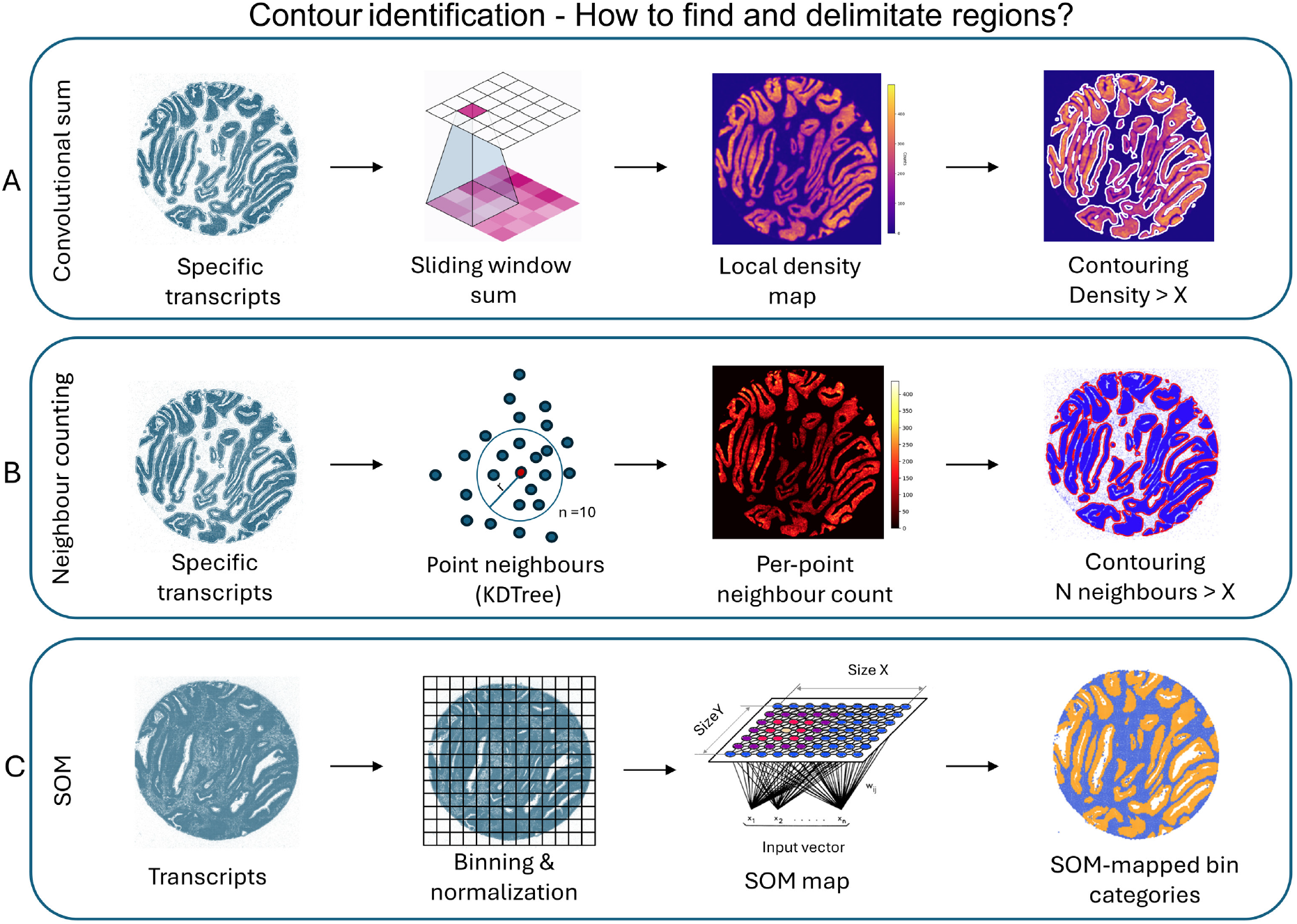
Contour identification strategies. A) Convolutional sum: A sliding window is applied across the spatial tissue coordinates, summing the expression counts of user-defined transcripts within each window. This generates a local density map where regions exceeding a defined threshold are contoured to highlight dense transcript areas. B) Neighbour counting: For each transcript, neighbouring points within a defined radius are counted using a KD-tree algorithm. This produces per-point neighbour counts, allowing contouring of regions where neighbour counts exceed a set threshold. C) SOM (Self-organizing map): Transcriptomic data is binned and normalized, then classified using an unsupervised SOM algorithm. The classified bins are mapped back, identifying spatial domains without requiring predefined sets of transcripts or thresholds.

In addition to convolutional-based region detection, GRIDGENE provides two alternative methods for mask generation: neighbourhood counting using KD-trees and unsupervised clustering via self-organizing maps (SOM). The neighbourhood counting method (Figure 2B), based on KD-tree algorithm (15), operates similarly to the convolutional sum approach but differs in implementation and output. Instead of applying a sliding window to create a continuous density map, this method uses a KD-tree to identify all neighbouring transcripts within a user-defined radius for each point. The result is a new set of point-wise neighbour counts, rather than a pixel-based image. However, because the method assigns values only to transcript locations (not the entire spatial grid), interpolation may be necessary to generate contiguous regions. Once interpolated, contours are extracted using density-based thresholding rules as with the convolutional approach. While this method may be computationally intensive for large-scale datasets, where the convolutional sum might be more efficient, it offers finer granularity for smaller, focused regions.

GRIDGENE also supports unsupervised tissue compartment identification through a SOM-based region detection method (Figure 2C). In this approach, transcriptomic data is first aggregated into spatial bins, which are then classified using the SOM algorithm (16–18), in an unsupervised fashion. The bin size determines the spatial resolution, while the number of SOM units defines how many distinct region types are identified. This method enables the discovery of spatially coherent expression domains without relying on predefined gene sets or thresholds, making it especially useful for exploratory analyses.

GRIDGENE provides key adjustable parameters — such as kernel size and minimum transcript counts — allowing users to fine-tune the granularity of detected regions. It also supports contour filtering, enabling the removal of regions based on customizable criteria (e.g., excluding contours below a certain size or with insufficient gene representation). For added flexibility, GRIDGENE allows users to manually define, and input contours generated through other approaches.

#### Tissue Mask Mapping and Expansion with GRIDGENE

GRIDGENE enables the conversion of contour data into tissue mask maps, providing options for refining and manipulating these masks:

1. From contours to masks: Upon receiving a set of contours, GRIDGENE generates corresponding masks that can be refined using various techniques:
  - Object filtering: removes small objects below a defined area threshold.
  - Morphological operations: applies operations such as gap filling and smoothing to improve mask quality and consistency.
  - Mask-level operations: combines, intersects, or differentiates masks to create meaningful tissue representations.
2. Expanding Masks: GRIDGENE supports mask expansion to explore the surroundings of defined regions. The expansions can be:
  - Simple distance-based expansion: expands masks uniformly by specified distances (e.g. 5 µm) from the mask’s edge.
  - Multiclass object analysis mode: ensures spatially distinct object expansions without overlap. Using Voronoi diagrams, GRIDGENE defines precise boundaries, making it ideal for comparing neighbourhoods of distinct objects that can be closely located.

#### Mask Information Retrieval

GRIDGENE provides comprehensive tools to extract data from masks:

- Transcript analysis: extracts transcript counts from masks.
- Morphological analysis: measures attributes such as area, perimeter, and shape descriptors.
- Hierarchy annotation: Identifies hierarchical and topological relationships between original objects and their expansions, linking derived regions to their parent contours.

This information can be retrieved over:

- Entire mask: measures across the full area covered by a type of mask (e.g., the full cancer or stroma compartment in a tumour).
- Individual objects: analyses each separate object within a mask type.
- Grids within objects: extracts data from regularly spaced grids placed inside each object.

#### Cell Segmentation Overlay

GRIDGENE outputs can be easily integrated with cell segmentation data by overlaying segmentation results onto tissue masks. It annotates cells based on their spatial relationships with defined regions, providing insights into cell distribution within specific regions.

### 2.2 Test case tumour microenvironment analysis

For carcinomas, the main biological compartments within a tumour tissue include epithelial regions, primarily composed of cancer cells, and stroma regions, which include stromal and immune cells. Pathologists typically identify these regions routinely based on tissue morphology. In ST experiments, these can be delineated based on areas with high expression of defined gene sets, bypassing the need for cell segmentation. Deriving masks for major biological compartments (e.g., epithelial and stroma regions) can facilitate downstream analyses targeting specific compartments or the interfaces between them.

#### 2.2.1 Definition of cancer-stroma in CRC on CosMx and Xenium

As an illustrative example, we applied GRIDGENE on data derived from the CosMx and Xenium ST platforms, which differ in resolution and targets, to generate cancer and stroma masks in CRC samples. To achieve this, convolutional kernels were used to delineate empty regions, with low overall density of genes, and cancer regions based on cancer-specific genes. After identifying contours for cancer and non-empty regions, mask maps of cancer and stroma compartments were obtained (Figure 3A-B). A cancer mask is defined by the cancer contours and a stroma mask is derived by negative selection, encompassing all regions that are neither cancer nor empty (Figure 3C). The stroma is defined through negative selection because its heterogeneous nature makes it challenging to identify a universal set of defining genes.

**Figure 3.**
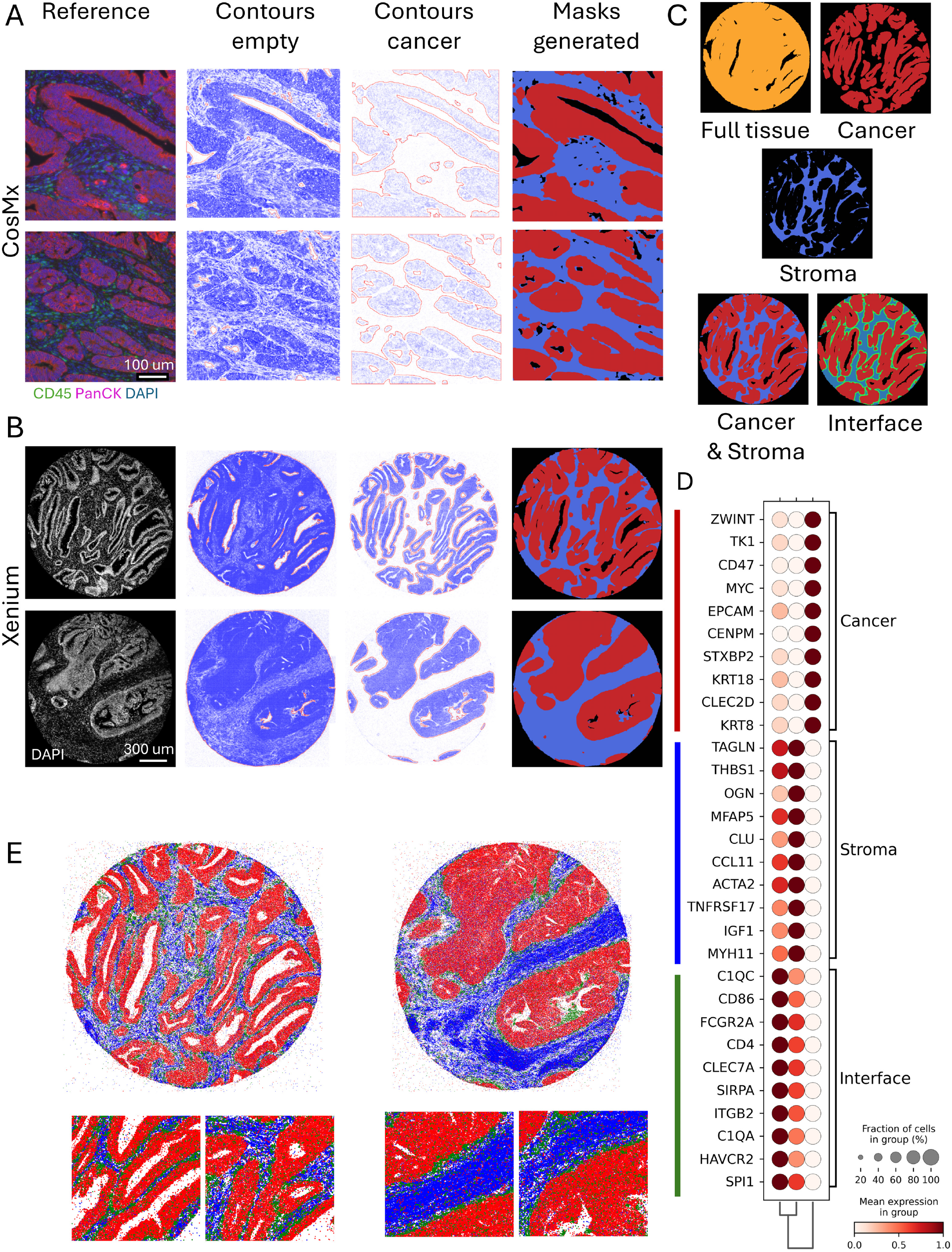
Cancer and stroma analysis with GRIDGENE. A) Example using CosMx data. Left to right: protein reference image showing Pan Cytokeratin (pink), nucleus (blue) and general membrane markers (green); contours identifying empty and cancer areas; and the final masks separating cancer and stroma regions. B) Example using Xenium data. Left to right: DAPI-stained nuclei reference image; contours identifying empty and cancer regions; and the generated masks for cancer and stroma areas. C) Schematic of the GRIDGENE pipeline to generate cancer stroma masks. First, all transcript points are used to identify tissue and empty regions. A cancer mask is defined using the cancer-specific transcripts, and stroma regions are determined by subtracting the cancer from the full tissue mask. The result is a complete division of the tissue into cancer and stroma compartments. To study the cancer-stroma interface, the cancer mask is further expanded into the stroma. D) For the Xenium cohort, cancer and stroma masks were defined, and 20 μm expansions of the cancer region were assessed. Transcript counts were retrieved from the masks, and differential gene expression across the three defined regions was calculated and visualized. E) Transcripts identified as differentially expressed at cancer stroma interface were plotted back onto two example cores, showing zoomed-in regions. Points are colour coded; red for cancer, blue for stroma and green for interface transcripts, highlighting the spatial localization of transcripts enriched in each mask..

Parameters of kernel size and density thresholds were adapted accordingly to each technology to reflect each platform’s unique characteristics. Furthermore, these parameters can also be adjusted to define regions with varying levels of granularity—ranging from highly detailed, cell-level distinctions to broader, smoother masks (Supplementary Figure 1).

#### 2.2.2 Cancer stroma interface analysis in Xenium CRC data

In addition to delineating cancer and stroma regions, it may be biologically relevant to characterise the interface between these compartments. To address this, the cancer mask can be expanded into the surrounding stroma areas for further analysis. We have defined the cancer and stroma areas as described in 2.2.1. The cancer mask was subsequently expanded by 10 and 20 µm into the stroma, and the transcript counts within these expanded regions were analysed. Differential gene expression across cancer, stroma, and interface masks was assessed (Figure 3D), and the differentially expressed transcripts were mapped back onto the tissues (Figure 3E), highlighting distinct transcriptomic profiles at the cancer-stroma interface. Of note, the same analysis can be applied independently to individual objects within a tissue, enabling differential assessment of their respective surrounding regions within the masks.

### 2.3 Single and Multiclass Object analysis

Beyond delineating major tissue compartments, defining regions based on pre-defined transcripts is especially valuable in cases where specific cell types cannot be reliably identified through cell segmentation and clustering.

#### 2.3.1 Definition of population-specific objects in CRC in CosMx

Although central to many ST workflows, cell segmentation presents several limitations. For instance, overlapping cells often pose significant challenges, particularly in heterogeneous tissues like tumour tissues. In addition, phenotyping based on clustering of segmented cells can be challenging, in particular for rare populations defined by a limited number of transcripts. Such cells are also more susceptible to misidentification due to transcript ‘contamination’ from neighbouring cells.

By employing the CosMx data, we sought to identify gamma delta (γδ) T cells, a rare immune cell subset that is otherwise difficult to detect through clustering and phenotyping of segmented cells (Figure 4A). To achieve this, we applied GRIDGENE’s convolutional approach to identify putative regions enriched for γδ T cell transcripts. Specifically, we identified areas where *TRGC* and *TRDC* transcripts, encoding for the T cell receptor (TCR) constant regions of *γδ* T cells, were within 5.5 µm of each other, with contours highlighting these regions. Beyond the presence of *TRGC* and *TRDC* transcripts in a confined area, specific criteria were applied, including thresholds on transcript counts, contour size, and transcript combinations, specifically requiring the sum of *TRGC* and *TRDC* counts to be higher than the sum of *TRAC* and *TRBC* (encoding for the TCR of *αβ* T cells), with at least one *TRGC* and one *TRDC* count per cell. The resulting regions represent areas potentially containing γδ T cells (Figure 4B) that can be subjected to further analysis.

**Figure 4.**
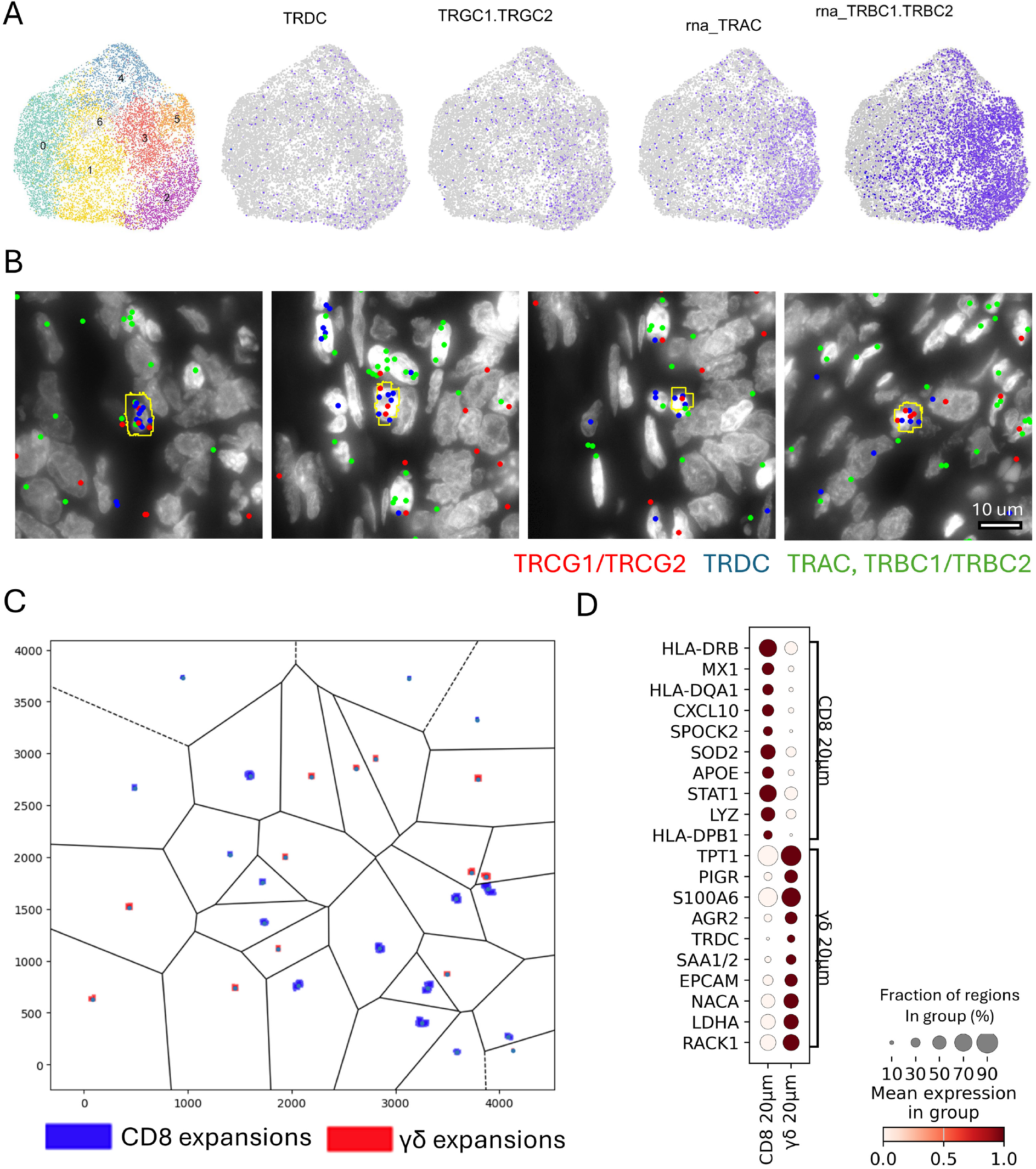
Population specific analysis with GRIDGENE in CosMx data. A) T-SNE of lymphocyte subclusters identified through cell segmentation. Expression of TRGC, TRDC, TRAC and TRBC transcript are projected onto the embedding, showing that γδ T cells cannot be distinguished from other T cells using segmentation and clustering alone. B) Representative tissue region showing GRIDGENE-identified γδ T cell objects (yellow boxes) overlaid on a DAPI-stained image. Transcript spots for TRGC1/TRGC2 (red), TRDC (blue), and TRAC/TRBC1/TRBC2 (green) are shown. γδ T cell objects are defined as regions with TRGC and TRDC transcript proximity (within 5.5 µm) meeting threshold criteria. C) Multiclass spatial analysis using a Voronoi diagram to partition a CRC sample (500 µm^2^ regions). γδ T cell expansions (red squares) and CD8+ T cell expansions (blue squares) are shown with non-overlapping boundaries. D) Differential expression analysis of transcripts within 20 µm from γδ and CD8+ T cell regions. The dotplot highlights genes with distinct expression in each context. Dot size represents the fraction of regions expressing the gene, and color intensity indicates average expression level.

#### 2.3.2 Multiclass Object analysis

Building on the principles of single-object analysis, this approach can be extended to simultaneously explore multiple objects. While single-object analysis can be repeated for different object types, it does not account for spatial overlaps or interactions between them, highlighting the need for a simultaneous, multi-object approach. The multiclass object analysis resolves this by generating Voronoi diagrams, which constrain expansions to areas closer to one object type than to any other. This ensures that regions around object type A are strictly associated with A and not influenced by proximity to object type B. We exemplify this approach using the previously defined γδ contours along with contours identifying CD8^+^ *αβ* T cell regions (Figure 4C). CD8^+^ T cell regions were defined using a kernel size of 5.5 µm to identify areas containing at least one transcript each of *CD8, TRAC*, and *TRBC*, where the combined *TRAC* and *TRBC* counts exceeded those of *TRDC* and *TRGC*, and *CD8* transcript levels were higher than *CD4*. To demonstrate the utility of this approach, we performed differential gene expression analysis between areas surrounding γδ T cell and CD8+ T cell regions, revealing distinct transcriptomic profiles in their respective microenvironments (Figure 4D).

### 2.4 Integration of GRIDGENE with cell segmentation

Defining regions of interest does not preclude the use of cell segmentation; rather, the two approaches can be easily integrated to facilitate specific tasks. GRIDGENE-generated masks can be overlaid onto cell segmentation data to assign spatial labels to cells—for instance, identifying them as intraepithelial (i.e., infiltrating epithelial compartments), stromal, or located at the interface of these compartments. Although cell segmentation approaches can yield similar results, they typically require definition of (continuous) neighbourhoods, which can be complex and less intuitive. By leveraging masks, these tasks become more straightforward and less time-consuming. In Figure 5, we illustrate an example of a sample analysed by Xenium ST technology overlaying the cancer, stroma, and interface masks with cell segmentation. We demonstrate that different immune cell types are differentially distributed across the different tissue compartments.

**Figure 5.**
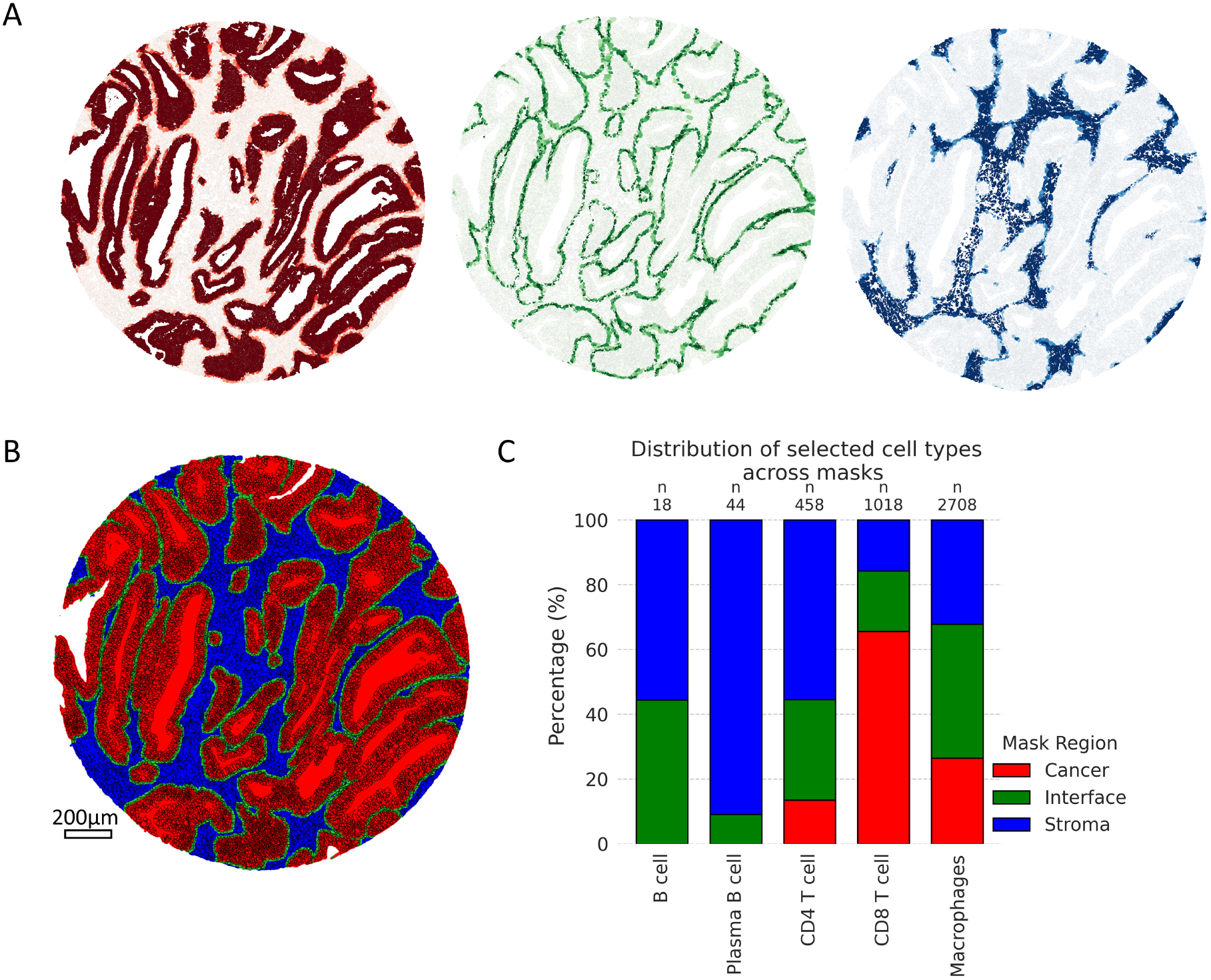
Overlay of Cell Segmentation and cancer-stroma-interface masks. A) Xenium core (1.5mm) with cell segmentation overlaid over cancer (red), stroma (blue), and interface (green) masks. B) Cell segmentation with cells coloured by the percentage of overlap with cancer (red), interface (green) and stroma (blue) masks. C) Distribution in percentage of specific immune cells across cancer, interface and stroma masks, revealing distinct spatial patterns: CD8 T cells are more concentrated in cancer regions, B cells are prevalent in the stroma and interface areas, and myeloid cells are more evenly distributed across all regions.

### 2.5 Alternative masking strategies

While this work primarily uses convolutional methods to derive contours, other masking approaches are also supported. In addition to convolutional techniques, GRIDGENE supports masks created using KDTree-based neighbour counting and Self-Organizing Maps (SOM) (Supplementary Figure 2).

Deriving from previous work in spatial proteomics (18), SOM-based mask generation applies the SOM algorithm (17) to bins of image data, classifying each bin in an unsupervised manner. The bin size can be adjusted to control the granularity of the masks, while the number of SOM units determines the number of resulting groups. In this study, we used SOM with two units, leading to an unsupervised separation of stroma and cancer compartments in Xenium data (Supplementary Figure 2B), distinction supported by differential gene expression observed between the two classes (Supplementary Figure 2C). This offers a possibility of deriving masks in an unsupervised fashion.

Neighbour counting based mask generation, in contrast to convolutional methods, relies on local neighbourhood point counts. The KD-tree algorithm, a space-partitioning structure, efficiently organizes points in multidimensional space and is commonly used to accelerate nearest-neighbour searches in local neighbourhood analysis. To exemplify this approach, we derived cancer and stroma regions in Xenium data (Supplementary Figure 2D) by calculating, for each point, the number of neighbouring transcripts within a 10 μm radius — both across the full set of genes and a cancer-specific gene set. Points with a high number of cancer gene neighbours were classified as cancer regions, while areas lacking nearby transcripts were designated as empty or stroma.

We also applied the KD-tree method to identify spatial associations between *TRGC* and *TRDC* transcripts (Supplementary Figure 2E) in order to identify *γδ* T cell regions in CosMx data. Specifically, for each TRGC or TRDC transcript, we searched for neighbouring TRDC or TRGC transcripts located within a 3 μm radius. This allowed us to identify transcripts that were spatially clustered, although the overall distribution remained sparse. Because such sparse clusters, often consisting of only a few transcripts, were insufficient for contour generation, we represented them by drawing circular masks that encompassed nearby transcripts. These proximity-based co-occurrences could also be projected onto segmented cell boundaries, enabling the identification of individual cells where *TRGC* and *TRDC* co-localize.

## 3. Discussion and Conclusions

Defining regions of interest in ST is essential for addressing specific biological questions. While cell segmentation-based methods are powerful, they may not always align with the needs of studies focused on tissue-level organization or areas defined by molecular density. For example, delineating cancer-stroma interfaces, identifying niches enriched for specific gene signatures, or exploring spatial relationships between regions of interest can be more straightforward when approached through region-level analysis. Regions defined by transcript density provide an intuitive framework for such tasks, reducing dependence on segmentation pipelines. GRIDGENE introduces a straightforward method to define and analyse regions based on transcript density. By leveraging user-defined gene sets and flexible kernel-based convolutional operations, the tool facilitates the identification of biologically meaningful areas without relying on cell segmentation. Its ability to generate cancer, stroma, and interface regions in tumour tissues was demonstrated effectively, and additional applications, such as identifying rare cell types, were also explored. Moreover, its capacity to integrate with cell segmentation pipelines ensures it can complement traditional workflows.

We would like to note that the reliability of this approach depends on several methodological parameters that require careful consideration. First, the accuracy of GRIDGENE relies heavily on the quality and representativeness of the selected genes. The identification of regions is constrained by the availability and specificity of transcripts, which may introduce biases or limit applicability in poorly annotated data. Second, the choice of kernel size and density thresholds, while offering flexibility, requires careful parameter tuning to balance precision and robustness. Mis-parameterization could result in either over-segmentation or under-segmentation of regions. Another consideration is that while GRIDGENE bypasses cell segmentation challenges, it does not replace the need for cell-level analysis in certain contexts. Questions involving detailed cellular phenotyping or fine-scale interactions remain better addressed by traditional cell segmentation approaches or a hybrid strategy.

Taking into account these important considerations, GRIDGENE offers a complementary perspective in spatial omics analysis, particularly for questions that benefit from region-level insights. GRIDGENE provides an efficient and adaptable framework to define and analyse these regions, enabling researchers to address specific biological questions while mitigating some of the challenges posed by traditional cell segmentation.

## 4. Methods

### 4.1 GRIDGENE implementation

GRIDGENE is implemented in Python 3.10 and leverages several scientific libraries, including NumPy, Pandas (19), Matplotlib (20), SciPy (15), Scikit-image (21), and OpenCV (22). It follows a modular design, enabling each module to function independently. For instance, externally derived masks or contours can be used as inputs without requiring GRIDGENE-generated data. To ensure transparency and traceability throughout the pipeline, structured logging is implemented. Additionally, the system supports extensive plotting at each pipeline stage for visual validation.

#### 4.1.1 Convolution-based contour generation

To identify regions of enriched transcript density, we implemented a convolution-based contour detection approach. The method begins by summing expression values across all transcripts at each pixel, producing a 2D transcript abundance map. A square or circular kernel is then applied to compute local transcript densities. Contours are extracted from this smoothed image by applying user-defined thresholds and detecting connected regions. Depending on the selected direction (“higher” or “lower”), either high- or low-density regions are detected. Spurious contours are removed based on a minimum area criterion and the presence of non-zero transcript counts in the original data. The method also includes automated validation to discard noise and regions with no biological signal, enhancing both accuracy and interpretability. The contour operations are based on OpenCV library.

#### 4.1.2 Neighbourhood counting-based contour generation

As an alternative, enriched transcript areas can be calculated directly with neighbouring transcript counts, a method based on KD-tree and query ball tree algorithm (Scipy implementation). This method proceeds in several steps. First, all foreground pixels (i.e., values equal to 1) are extracted from a binary array and interpreted as spatial coordinates. These are indexed using a KD-tree structure to enable efficient local neighbour queries. Points are then filtered to retain only those that have at least *N* neighbouring points within a given radius, thus removing isolated detections and reducing noise.

From here, contours can be generated as described above. As transcripts are often sparse, to achieve better contours, OpenCV inpaint function can be used to interpolate values at missing positions. Alternatively, the filtered points can be grouped into spatial clusters using the DBSCAN algorithm (23) (scikit-learn (24) implementation) and contours can be generated from these labelled groups of points. If the number of annotated points is very small, simpler methods of deriving contours are available: the simplest method computes a minimum enclosing circle for each cluster, ensuring a compact representation of local signal accumulation. Alternatively, convex or concave hulls (alpha shapes) are computed around each point cloud, depending on the desired boundary fidelity.

#### 4.1.3 SOM-based contour generation

This method uses grid binning based on spatial coordinates, with customizable grid shapes, and leverages the AnnData (25) structure for preprocessing using Scanpy (26). Spatial bins are classified using a SOM (17) implemented via MiniSOM (16). The classification results are mapped back to the original coordinates to define regions of interest. Statistical validation of SOM-based separation is available through t-tests, providing an assessment of the significance of spatial clustering.

#### 4.1.4 Mask generation

From a set of contours binary masks are generated using OpenCV. Several post-processing operations are implemented, including area filtering to remove small, disconnected components; gap filling; and morphological operations (erosion, dilation, opening, closing) to refine boundaries and reduce noise. Mask subtraction is also supported, allowing specific regions to be isolated by subtracting one or more masks from a base mask.

Masks can be expanded via OpenCV dilation and can be constrained by other masks (e.g., expanding a tumour mask only within the stroma). To avoid overlaps in multiclass mask expansions, a Voronoi-based constraint system is implemented. Using centroids of all objects, a Voronoi diagram is generated (via SciPy), and each object’s expansion is limited to its respective Voronoi cell, ensuring that expansion remains closest to its source object. Dedicated classes support the integration and expansion of multiple masks into cohesive, non-overlapping tissue masks.

#### 4.1.5 Mask information retrieving

All mask objects are labelled using scikit-image’s labelling tools. Transcript counts can be retrieved for the entire mask, individual labelled objects and gridded subregions within objects. This process supports multiprocessing for improved performance on large datasets. Morphological characteristics (e.g., area, perimeter, shape descriptors) are computed using scikit-image regionprops function. A hierarchical analysis is also supported: expanded objects can be traced back to their originating (“parent”) object by storing and mapping object labels throughout the pipeline. The information is stored in a pandas’ DataFrame.

#### 4.1.6 Cell segmentation overlay

Cell segmentations (in mask or polygon format) can be overlaid onto the generated masks. For each cell, the number of overlapping pixels with each mask is computed, and the percentage of coverage is reported.

### 4.2 Data

Two CRC cohorts were analysed: one consisting of 20 regions of interest, profiled using CosMx (5), and another with 20 regions of interest, profiled with Xenium (4). In the CosMx dataset, regions of interest are square (500 *μ*m × 500 *μ*m), corresponding to images of 4245 × 4245 pixels (approximately 0.12 *μ*m per pixel). CosMx data includes measurements for 999 genes from a standard panel. In contrast, Xenium tissue cores are circular, 1500 *μ*m in diameter, corresponding to 1500 pixels (1 *μ*m per pixel), and are profiled for 480 genes using a custom panel. These differences in resolution, pixel size, and gene coverage necessitate platform-specific analytical strategies and parameter adjustments to ensure accurate region delineation and to account for the unique strengths of each platform.

### 4.3 Cancer-stroma definition in CosMx and Xenium

First, tissue-containing regions were distinguished from empty areas. For CosMx, empty regions were identified using a kernel size of 80 × 80 pixels (approximately 10 × 10μm), with areas containing fewer than 140 transcript counts classified as empty. For Xenium, a 10 x 10 *μ*m kernel was used, and regions with fewer than 40 detected transcripts were considered empty. Next, cancer regions were identified. In CosMx, this was done using a cancer-specific transcript set and defining objects with a kernel of 80 × 80 pixels (approximately 10×10μm) with at least 40 of cancer-specific transcript counts. For Xenium, a kernel of 10×10μm, and objects were retained if they contained at least 10 transcript of the cancer transcripts. Cancer transcripts were chosen based on prior knowledge: CosMx cancer transcripts were *EPCAM, KRT19, KRT8, KRT18, KRT17, CEACAM6, SPINK1, CD24, S100A6, RPL37, S100P*. For Xenium the transcripts used were *EPCAM, SMIM22, CLDN3, KRT18, LGALS4, KRT8, ELF3, TSPAN8, STMN1, CD47, MYC, LGALS3*. The differences in these parameters reflect the inherent variations in resolution and transcript coverage between the two technologies, highlighting the importance of tailoring analyses to each platform’s characteristics.

After identifying contours for cancer and non-empty regions, mask maps of cancer-stroma were derived. A cancer mask is defined by the cancer contours and a stroma mask is derived by negative selection, encompassing all regions that are neither tumour nor empty.

### 4.4 Cancer-stroma interface analysis

For the Xenium cohort, cancer tissue masks were expanded by 20 µm into the adjacent stroma, and transcript counts within these expanded regions were analysed. Transcript data from three distinct regions—the original cancer mask, the 20 µm peri-tumoral expansion, and the remaining stroma— were extracted for each sample. Transcript counts were normalized to a target sum of 1e-4 and log-transformed. Principal component analysis (PCA), neighbourhood graph computation, and UMAP embedding were performed using functions from the Scanpy package (26). Gene expression differences across the three regions were ranked using the Wilcoxon test, considering only genes with a minimum fold change of 1. Expression values for stroma, cancer, and their interface were visualized in two representative tissue cores.

### 4.5 Population specific objects definition

Using the CosMx cohort, we identified putative γδ T cell regions via a convolutional sum approach, applying a 45 × 45-pixel kernel (approximately 5.5 μm) and requiring at least two transcripts from *TRGC* and/or *TRDC*. These regions were further filtered to include only those containing at least one *TRGC* and one *TRDC* transcript, with the combined *TRGC* + *TRDC* count exceeding the total *TRAC* + *TRBC* count.

CD8+ T cell regions were identified using the same 45 × 45-pixel kernel, requiring a minimum of two transcripts from the set {*TRAC, TRBC1*/*TRBC2, CD8A, CD8B*}. Regions were retained only if they included at least one *CD8* (A or B) transcript, one TRAC, and one *TRBC* transcript. Additionally, the combined *TRAC* + *TRBC* count had to exceed that of *TRDC* + *TRGC*, and CD8 transcript levels had to be greater than *CD4*.

Both γδ T cell and CD8+ T cell regions were converted into binary masks. To generate spatial expansions, we applied multiclass object analysis using Voronoi diagrams to constrain each expanded region to the nearest cell type. A 20 μm expansion was generated for both γδ and CD8+ T cell regions, and these were analysed similarly to cancer–stroma interface masks, including normalization, principal component analysis (PCA), and differential gene expression (DGE) analysis. DGE was performed directly between the expanded γδ and CD8+ T cell regions to compare their surrounding transcriptomic environments.

### 4.6 Cell segmentation

Cell segmentation for the CosMx cohort was provided by the manufacturer as previously described (5). For the Xenium cohort, we improved segmentation quality by applying Baysor (8), using the default Xenium segmentation as a prior with a confidence level of 0.7. Baysor parameters were set as follows: a minimum of 30 molecules per cell, a spatial scale of 10, and three clusters. All other parameters were left at their default values.

### 4.7 Cell phenotyping

Similar cell phenotyping approaches were applied to both the Xenium and CosMx cohorts. Cell counts were extracted, and low-transcript cells were excluded. The data were log-normalized, followed by computation of nearest neighbours, PCA, and UMAP. Clustering was performed using the Leiden algorithm. Initially, stromal and cancer clusters were separated. The stromal cluster was then further refined to identify general stromal cell types, including fibroblasts and immune cells such as B cells, T cells, and myeloid cells.

### 4.8 Cell segmentation and mask overlay

Cancer-stroma interface masks were overlaid with the corresponding cell segmentation and phenotyping data. For each segmented cell, the proportion of its area overlapping with each mask was calculated. Cells were then assigned to the mask covering the largest portion of their area, and the frequencies of T cells, B cells, and myeloid cells were computed for each mask category

### 4.9 SOM-based masks

For the cancer-stroma analysis in the Xenium cohort, an alternative approach based on SOM was applied. Images were divided into 5×5 μm bins, and bins containing fewer than 10 transcripts were excluded. Transcript counts per bin were log-normalized, and a SOM with a (2,1) shape was trained for 5,000 iterations using a sigma of 0.5 and a learning rate of 0.5. This configuration classified the bins into two distinct categories. To assess whether these categories corresponded to stromal and epithelial compartments, a t-test was performed to rank the top genes associated with each class. The classified bins were then mapped back onto the tissue samples, and masks were generated accordingly.

### 4.10 Neighbour counting based masks

For exemplification purposes, we use KD-tree-based mask generation to also derive cancer and stroma masks in Xenium. A KD-tree (implemented from Scipy (15)) was constructed using the spatial coordinates of the transcriptomic points to enable efficient nearest-neighbour searches in Euclidean space. For each point in the dataset, the query_ball_point method was employed to retrieve the indices of all neighbouring points within a 10-pixel radius, enabling the computation of local neighbourhood densities with optimized performance for large-scale spatial data. Given the inherent sparsity of transcriptomic points, we applied inpainting using the OpenCV (22) inpaint function to interpolate missing values and optimize contour detection. From this smoothed neighbourhood map, contours were generated following the same procedure described in 4.5.

Additionally, we used local point search (based on KD-tree) to identify the points of *TRGC* and *TRDC* close to each other using a radius of 25 pixels in CosMx data. We label all the sets of close points as belonging to the same object using DBSCAN (23) a Density-Based Spatial Clustering implemented using scikit-learn (24). Because only these transcripts were classified and the density of points was insufficient to generate contours, we instead drew circular regions that enclosed the identified points. These regions were then filtered using the same exclusion criteria described in Section 4.2.4.

## 5. Data and code availability

The GRIDGENE pipeline is fully available and ready to use at https://github.com/deMirandaLab/GRIDGENE. The repository also contains all the necessary code to replicate the methods comparison, along with detailed Jupyter notebooks for easy understanding and implementation.

## 6. Acknowledgments

A.M.S. was supported by the Portuguese Foundation for Science and Technology-FCT and FSE through PhD fellowship:2020.0786BD. N.F.C.C.d.M. is funded by the European Research Council under the European Union’s Horizon 2020 Research and Innovation Programme (grant agreement no. 852832)) and by VIDI ZonMW (project number 09150172110092).

The authors declare no conflicts of interests.

