## Supplementary Materials for "GRIDGENE: Guided Region Identification based on Density of GENEs – a transcript density-based approach to characterize tissues by spatial transcriptomics"

### Supplementary information

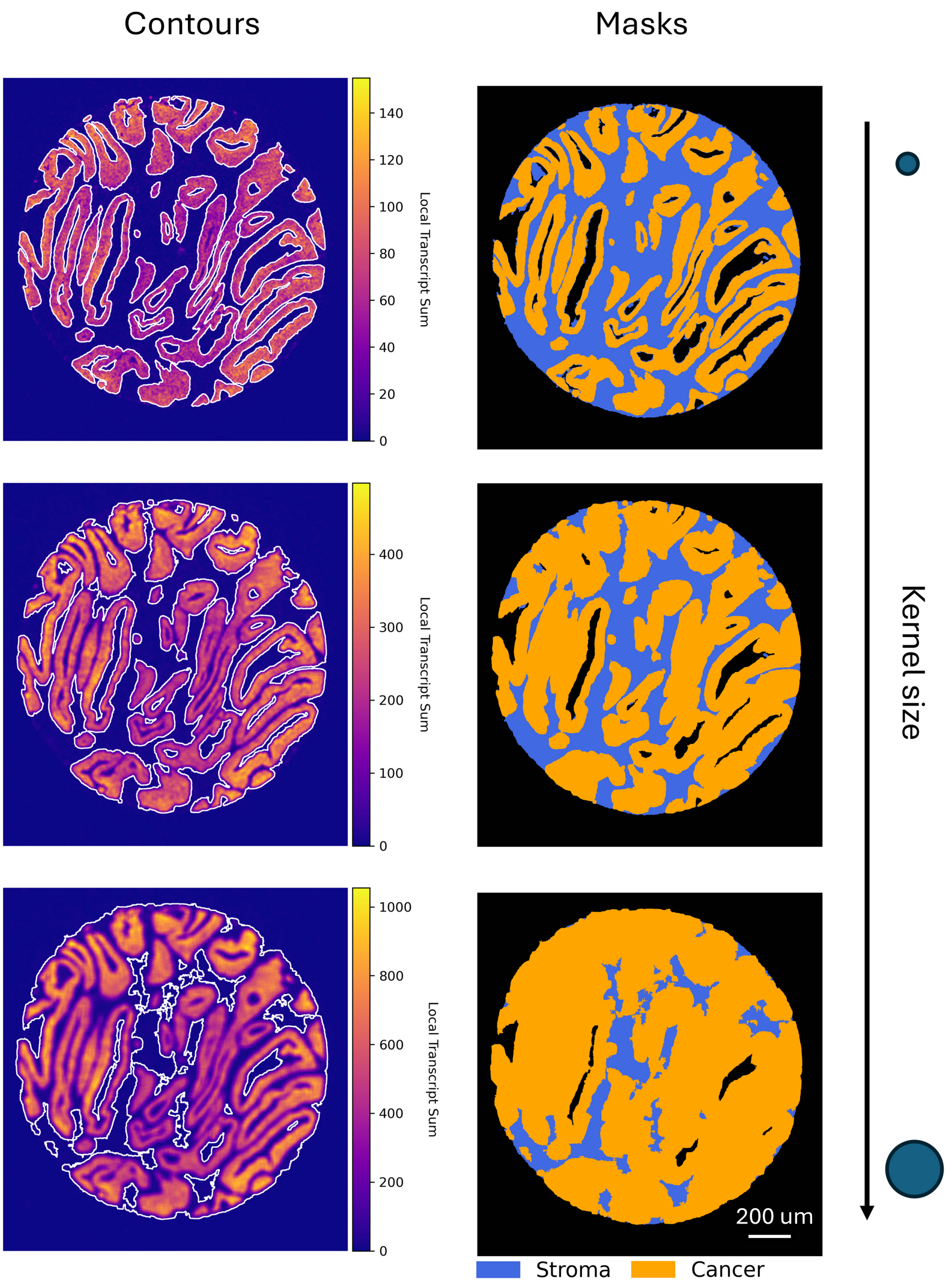


*Supplementary figure 1 – Influence of Kernel Size on Mask Generation. Example from a Xenium dataset illustrating how varying the kernel size affects the shape and granularity of cancer masks. The stroma mask remains constant, while the cancer mask is computed using kernel sizes of 10 µm, 20 µm, and 30 µm for the convolutional sum. Larger kernel sizes produce smoother and more generalized contours, while smaller kernels result in more fine-grained and detailed masks, potentially capturing localized structures.*

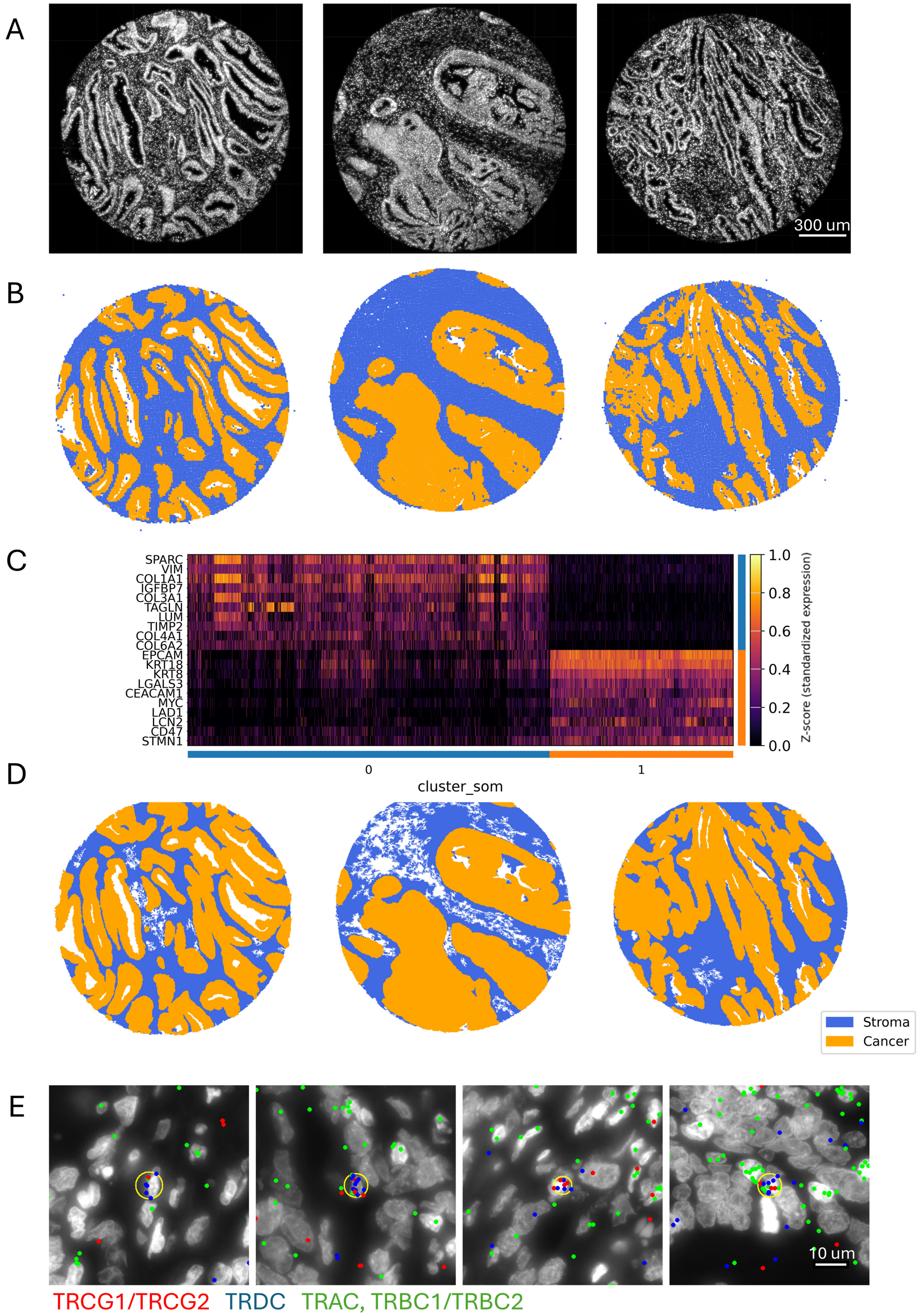


*Supplementary figure 2 – Alternative strategies for spatial masking using GRIDGENE. (A) Xenium DAPI references from three CRC samples. (B) Cancer and stroma regions were masked using a self-organizing map (SOM)–based approach, where 10×10 pixel grids were clustered using a SOM with two units. Although unsupervised, this method appears to separate stroma and cancer compartments. (C) Heatmap showing genes differentially expressed between the two SOM-derived clusters, supporting their identification as stroma and cancer. (D) Alternative cancer/stroma masks were*

*generated by calculating local point density using a KD-Tree with a 10 μm radius, then deriving contours based on neighbourhood counts. (E) Putative γδ T cells were identified by locating TRGC and TRDC transcripts that are ≤3 μm apart. Circular regions were drawn around these pairs and discarded if TRAC or TRBC signals were more abundant. These regions can be overlaid with cell segmentation data to identify γδ T cells or used directly as spatial regions of interest.*
